# Chaperone mRNA secondary structure does not strongly track thermal adaptation in bacteria, but *groL* coding sequence shortens in its disordered C-terminal tail

**DOI:** 10.64898/2026.11.10.765738

**Authors:** Sudheesh Singh, Vaibhav Sahu, Sushree Sradha, Vedikaa Dhiman

**Affiliations:** National Institute of Technology, Raipur; Manipal University Jaipur

**Keywords:** mRNA secondary structure, thermal adaptation, GroEL, chaperonin, phylogenetic comparative methods, RNAfold

## Abstract

Ribosomal and transfer RNAs show clear compositional and structural signatures of adaptation to growth temperature, but messenger RNA does not, and the generality of this asymmetry at individual loci remains untested. We asked whether the two principal bacterial chaperone transcripts, dnaK and groL, encode systematically different mRNA secondary structures in thermophiles relative to mesophiles. Using a phylum-matched cohort of 50 bacterial strains (25 mesophiles, 25 thermophiles; balanced across Actinobacteria, Bacteroidetes, Firmicutes and Proteobacteria), we folded each coding sequence at 37°C and 55°C and compared fifteen thermodynamic and structural descriptors per gene. Raw folding metrics showed no significant group differences after multiple-testing correction. However, after controlling for dinucleotide composition with Altschul-Erickson shuffles, thermophile sequences, particularly groL, retained modestly more residual secondary structure than mesophile sequences. The strongest and cleanest signal was not structural: groL coding sequences are shorter in thermophiles (mean 1626.5 versus 1636.3 nt; Cliff’s δ = 0.571, P = 5.1×10^-4^), a relationship that scales continuously with optimal growth temperature and persists after phylogenetic generalised least squares correction (n = 50, Pagel’s λ = 0.997, slope = -0.356 nt°C^-1^, P = 6.0 × 10^-5^). Protein-level alignment localised this difference: approximately three-quarters of the excess deletion signal falls within the terminal ∼25 residues corresponding to the intrinsically disordered, glycine/methionine-rich C-terminal tail of GroEL. Thermal adaptation in these loci therefore appears to act primarily on a disordered peptide extension, with only a modest residual signature detectable in mRNA folding after composition is controlled.

## Introduction

Temperature shapes macromolecular sequence and structure across the tree of life, and the search for molecular signatures of thermal adaptation has a long history. At the RNA level, this search has produced an informative asymmetry. Transfer RNA and ribosomal RNA show strong positive correlations between guanine-cytosine content and optimal growth temperature (T_opt_), and rRNA secondary structure content increases with T_opt_ in a manner not explained by nucleotide composition alone. Messenger RNA behaves differently. In the most comprehensive analysis to date, Jegousse *et al*.^1^ examined 9,624 mRNA sequences drawn from 172 essential protein-coding genes across 287 bacterial species spanning T_opt_ from 4°C to 95°C, and found no correlation between mRNA GC content and T_opt_, a *negative* correlation between predicted paired-nucleotide fraction and T_opt_, and critically no significant difference between natural mRNA sequences and dinucleotide-matched random controls. They concluded that mRNA is compositionally “pre-wired” for flexibility rather than structurally optimised for its host’s thermal niche.

Against this, work in eukaryotes has found genuine thermal signals in mRNA folding energetics. Liao *et al*.^2^ showed that the free energy change of secondary structure ensemble formation (ΔG_fold_) for cytosolic malate dehydrogenase mRNA correlates significantly with adaptation temperature across 25 marine mollusc species, and attributed this to synonymous G+C substitutions that tune structural stability without altering the encoded protein. Whether such locus-specific mRNA thermal tuning occurs in bacteria and specifically in genes whose function is itself thermal, is not resolved by either study.

Chaperone genes are the natural test case. The Hsp70 (DnaK) and Hsp60 (GroEL/GroES) systems carry the majority of the cellular protein-folding load, and chaperone-mediated proteome reallocation is a principal determinant of bacterial growth thermosensitivity^3^. If any transcripts were under selection to fold appropriately at their host’s operating temperature, transcripts encoding the heat-shock machinery would be plausible candidates.

We note at the outset that our two genes are not equally novel territory. Gil *et al*.^4^, the minimal-gene-set analysis from which Jegousse *et al*. drew their gene list, explicitly identifies GroEL and GroES as the best-conserved chaperones and as essential in *Escherichia coli* and *Bacillus subtilis*, so *groL* is likely already represented within that earlier dataset. The same analysis describes the DnaK-DnaJ-GrpE system as non-essential in free-living bacteria, so *dnaK* is probably not. Our contribution is therefore not the first examination of *groL* mRNA structure but a differently designed one: we use explicit phylum matching rather than T_opt_-binned averaging across an unselected species set, we fold at two temperatures rather than one, we apply phylogenetic correction, and we ask where within a gene any detected difference is located. This last question, to our knowledge, has not previously been addressed for a bacterial chaperonin.

## Materials and methods

### Strain selection

Fifty bacterial strains were selected to give 25 mesophiles and 25 thermophiles matched by phylum: Actinobacteria (6 + 6), Bacteroidetes (6 + 6), Firmicutes (7 + 7) and Proteobacteria (6 + 6). Matching was imposed by design so that thermal-group contrasts would not be confounded with clade membership. Optimal, minimum and maximum growth temperatures were taken from the TEMPURA database^5^. Mesophiles spanned T_opt_ 29-44°C and thermophiles 50-78°C; hyperthermophiles (T_opt_>80°C) were excluded by design so that the folding temperatures used remained physiologically interpretable for both groups. Archaea were excluded. Coding sequences for *dnaK* and *groL* were retrieved from NCBI by locus tag, one orthologue per strain.

### RNA secondary structure prediction

Each coding sequence was folded with RNAfold from the ViennaRNA package^6^, which computes the minimum free energy structure by the algorithm of Zuker and Stiegler^7^ and equilibrium base-pairing probabilities from the McCaskill partition function^8^. Folding was performed at 37°C and at 55°C using default parameters otherwise. For each sequence and temperature we recorded minimum free energy (MFE), MFE normalised per nucleotide, ensemble free energy, ensemble diversity, the MFE-to-ensemble gap, the MFE-to-centroid gap, and the fraction of paired bases in the MFE structure. GC content and coding sequence length were recorded once per sequence. ΔΔG was computed as MFE(55°C) - MFE(37°C). Ensemble diversity was normalised per nucleotide to remove the dependence of this quantity on transcript length.

### Dinucleotide-preserving shuffle controls

To test whether any residual secondary-structure signal remained after accounting for nucleotide composition, each coding sequence was subjected to dinucleotide-preserving randomisation using the algorithm of Altschul and Erickson^15^. This procedure generates sequences that exactly preserve both mononucleotide and dinucleotide frequencies (and therefore the same GC content and nearest-neighbour stacking potential) while destroying higher-order sequence order. For each native sequence we generated 100 independent shuffled versions. All shuffled sequences were folded with RNAfold under identical conditions (37 °C and 55 °C) as the corresponding native sequence. A structural z-score was then calculated for every native sequence as:

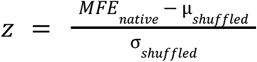

where μ_shuffled_ and σ_shuffled_ are the mean and standard deviation of the minimum free energies of the 100 shuffled sequences. Negative z-scores indicate that the native sequence is more stably folded than expected from its dinucleotide composition alone. Robustness of the z-score estimates was confirmed by repeating the entire procedure with 20 and with 50 shuffles; the group-level statistics remained essentially unchanged. Z-scores were compared between mesophiles and thermophiles by two-sided Mann-Whitney U tests and were correlated with T_opt_ by Spearman rank correlation, exactly as for the primary thermodynamic descriptors.

### Statistical analysis

Fifteen descriptors per gene were compared between thermal groups using a two-sided Mann-Whitney *U* test, applied uniformly to every descriptor. No alternative normalisations or one-sided variants were substituted for individual comparisons. Effect sizes are reported as Cliff’s δ^9^. The four composition-controlled z-score comparisons were analysed with the same test. All 34 *P*-values (30 primary descriptors + 4 z-scores) were corrected together by the Benjamini-Hochberg procedure^10^ at a false discovery rate of 5%; Bonferroni-corrected values are also given. Continuous relationships with T_opt_ were assessed by Spearman rank correlation.

### Phylogenetic correction

16S rRNA gene sequences were retrieved for all 50 strains, aligned with MAFFT v7 under the `--auto` strategy^11^, and a maximum-likelihood tree was inferred with FastTree 2 under a generalised time-reversible model with CAT rate categories^12^. A Brownian-motion phylogenetic variance-covariance matrix was constructed from shared root-to-most-recent-common-ancestor path lengths, and Pagel’s λ transformation was applied to the off-diagonal elements^13^. Phylogenetic generalised least squares models of coding sequence length against T_opt_ were fitted by generalised least squares^14^ with λ estimated by maximum likelihood; the likelihood surface was evaluated on a grid over λ ∈ [0, 1] to confirm that the optimum was interior or a genuine boundary maximum rather than a numerical artefact. Because no fossil calibration exists for this strain set, branch lengths are in expected substitutions per site and the tree is treated as a non-clock phylogeny; this is standard practice for microbial comparative analyses in the absence of a molecular clock but is an assumption of the model rather than a property of the data.

### Localisation of length differences

All 50 GroEL protein sequences were aligned with MAFFT v7. For each alignment column we computed the fraction of mesophile and of thermophile sequences carrying a gap, and took the difference (thermophile minus mesophile) as the excess deletion rate. Columns were mapped to *E. coli* GroEL residue numbering (548 residues) using the gap-free positions of the *E. coli* sequence as an index.

### Data and code availability

All source data, sequence alignments, the 16S rRNA phylogenetic tree, and analysis scripts supporting this study are publicly available at https://github.com/Sudheesh02/chaperone-mrna-thermal-adaptation.

## Results

### Structural and thermodynamic descriptors do not separate the thermal groups

Of 34 pre-specified comparisons (30 primary descriptors + 4 composition-controlled z-scores), six reached nominal significance at P < 0.05. Two survived Benjamini-Hochberg correction at a 5% FDR (Figure 1): *groL* transcript length and the groL structural z-score at 55°C. Both of these also survived Bonferroni correction. Among the primary thermodynamic descriptors, no minimum free energy measure, in either raw or per-nucleotide form, at either folding temperature, in either gene, differed detectably between mesophiles and thermophiles (all P ≥ 0.38). Neither did GC content (*dnaK* P = 0.31; *groL* P = 0.21), paired-base fraction, ensemble gap, or ΔΔG between the two folding temperatures. Ensemble diversity per nucleotide at 55°C reached P = 0.030 for *groL* and P = 0.065 for *dnaK*, neither surviving correction. At the level of raw mRNA folding thermodynamics, then, these two chaperone transcripts are indistinguishable between thermal niches, a result consistent with, and extending to single-locus resolution, the genome-wide conclusion of Jegousse *et al*.^1^

**Figure 1.**
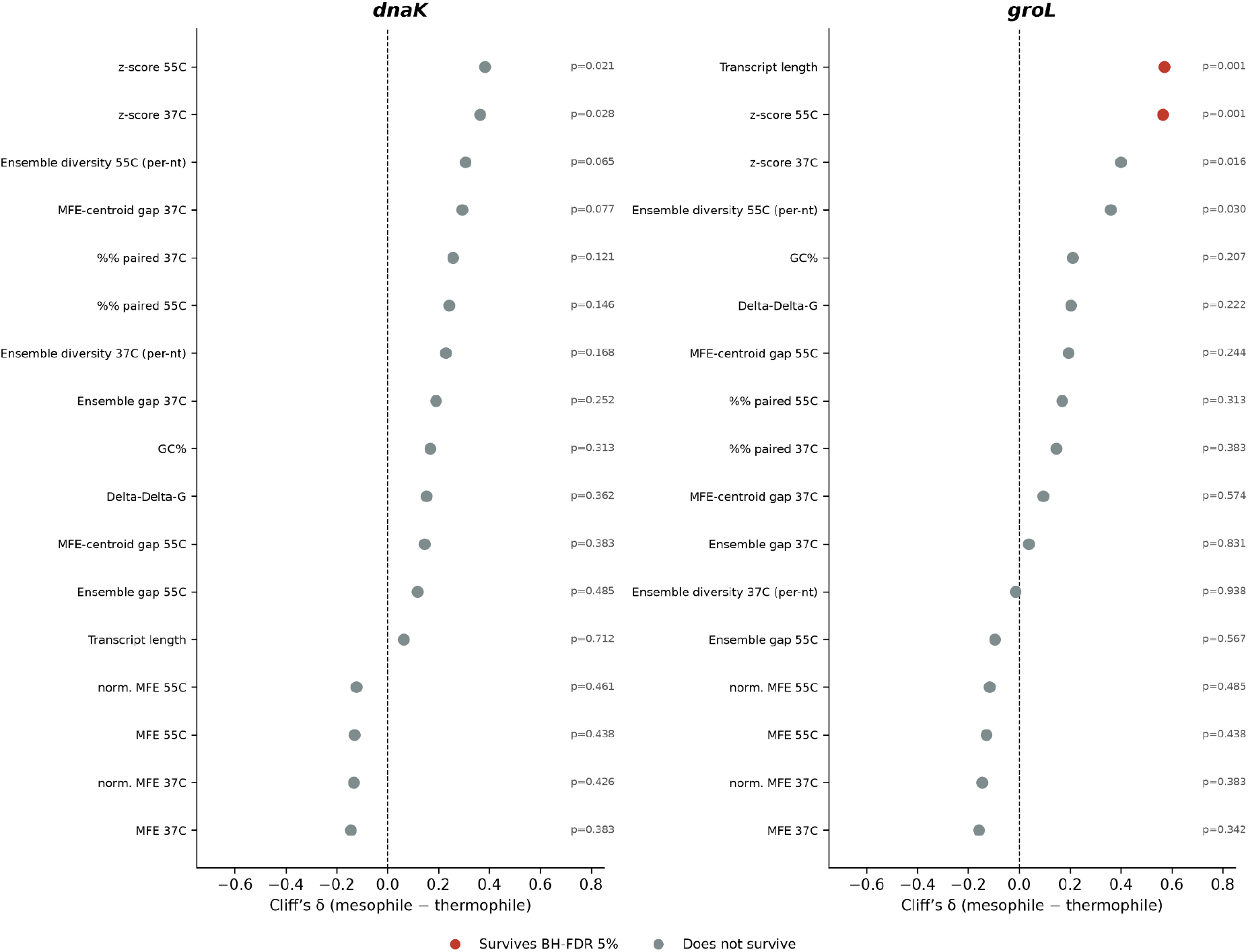
Effect sizes and significance for all 34 pre-specified comparisons between mesophiles and thermophiles, shown separately for *dnaK* (left) and *groL* (right). Points give Cliff’s δ (mesophile - thermophile); *P*-values are from two-sided MannWhitney *U* tests applied uniformly to every descriptor. Red indicates the two comparisons surviving Benjamini-Hochberg correction at a 5% false discovery rate; grey indicates all others. n = 25 per group per gene.

### Composition-controlled structural z-scores reveal residual signal

Although the primary thermodynamic descriptors showed no group difference, a residual structural signal became apparent once dinucleotide composition was controlled. For each coding sequence we generated 100 dinucleotide-preserving shuffles, folded them under the same conditions as the native sequence, and calculated a z-score measuring how much more (or less) stable the native fold was relative to the composition-matched null. Thermophile *groL* sequences yielded significantly more negative structural z-scores than mesophile sequences at 55°C (mean z = -1.41 versus -2.77; Mann-Whitney *P* = 6.4 × 10^-4^, Cliff’s δ = +0.56). This comparison remained significant after Benjamini–Hochberg correction across the full set of 34 tests (30 primary descriptors + 4 z-score comparisons). The corresponding difference at 37°C (*P* = 0.016) and the parallel tests for *dnaK* did not survive multiple-testing correction.

Results were essentially unchanged when the analysis was repeated with 20 or with 50 shuffles. Thus, while raw folding free energies do not differ between thermal groups, thermophile *groL* transcripts retain modestly more secondary structure than expected from their dinucleotide composition alone at the higher folding temperature.

Mean z-scores were calculated from 100 Altschul-Erickson dinucleotide-preserving shuffles per sequence. Negative values indicate that the native sequence is more stably folded than expected from its dinucleotide composition alone. P-values are from two-sided Mann-Whitney U tests. Cliff’s δ is the effect size (positive values indicate more negative z in thermophiles).

**Table 1.**
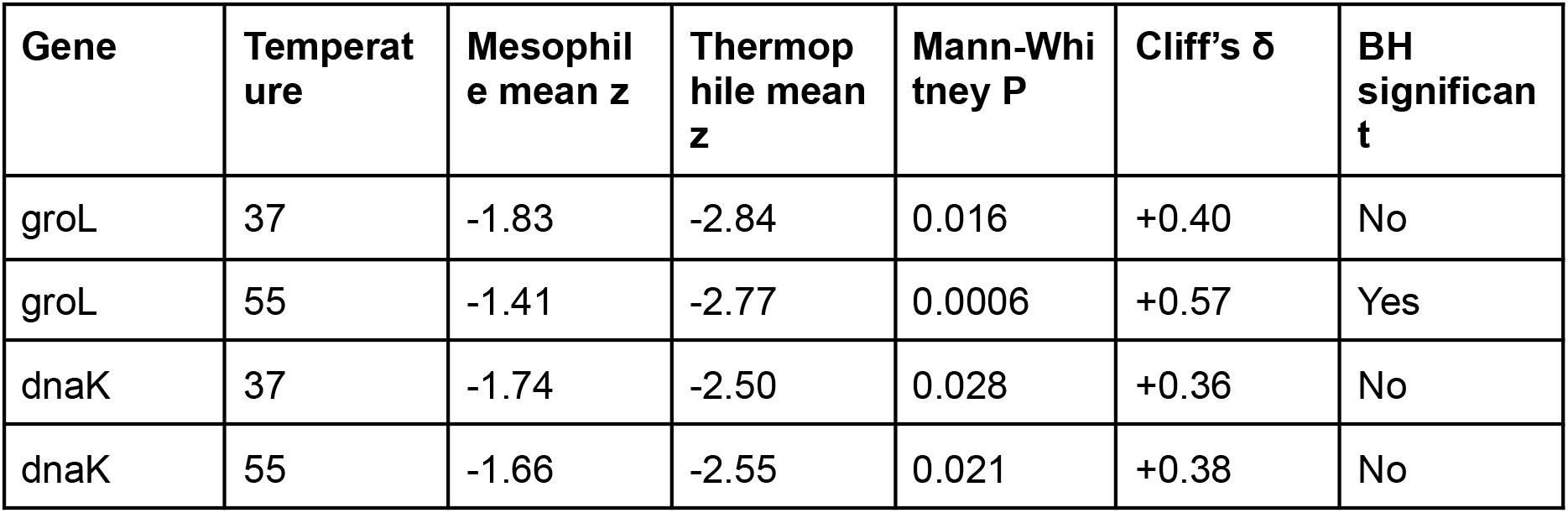
Dinucleotide-controlled structural z-scores for groL and dnaK.

| Gene | Temperature | Mesophile mean z | Thermophile mean z | Mann-Whitney P | Cliff’s δ | BH significant |
|---|---|---|---|---|---|---|
| groL | 37 | -1.83 | -2.84 | 0.016 | +0.40 | No |
| groL | 55 | -1.41 | -2.77 | 0.0006 | +0.57 | Yes |
| dnaK | 37 | -1.74 | -2.50 | 0.028 | +0.36 | No |
| dnaK | 55 | -1.66 | -2.55 | 0.021 | +0.38 | No |

### The one robust difference is coding sequence length, and only in *groL*

Mesophile *groL* coding sequences average 1636.3 ± 10.1 nt and thermophile sequences 1626.5 ± 7.2 nt, a difference of 9.8 nt or approximately three codons (Cliff’s δ = 0.571, *P* = 5.1×10^-4^; Bonferroni-corrected *P* = 0.017). The effect is not a threshold phenomenon at the mesophile/thermophile boundary but scales continuously with Topt (Spearman ρ = -0.589, *P* = 6.8 × 10^-6^; Figure 2). It is directionally consistent within each of the four phyla examined separately (Actinobacteria ρ = -0.790, *P* = 0.0022; Bacteroidetes ρ = -0.710, *P* = 0.010; Firmicutes ρ = -0.657, *P* = 0.011; Proteobacteria ρ = -0.820, *P* = 0.001), indicating that the pattern is not driven by any single clade. *dnaK* shows no comparable relationship (mean difference -3.1 nt; ρ = -0.057, *P* = 0.70), so the effect is specific to *groL* rather than reflecting a general tendency for thermophile genes to be shorter.

**Figure 2.**
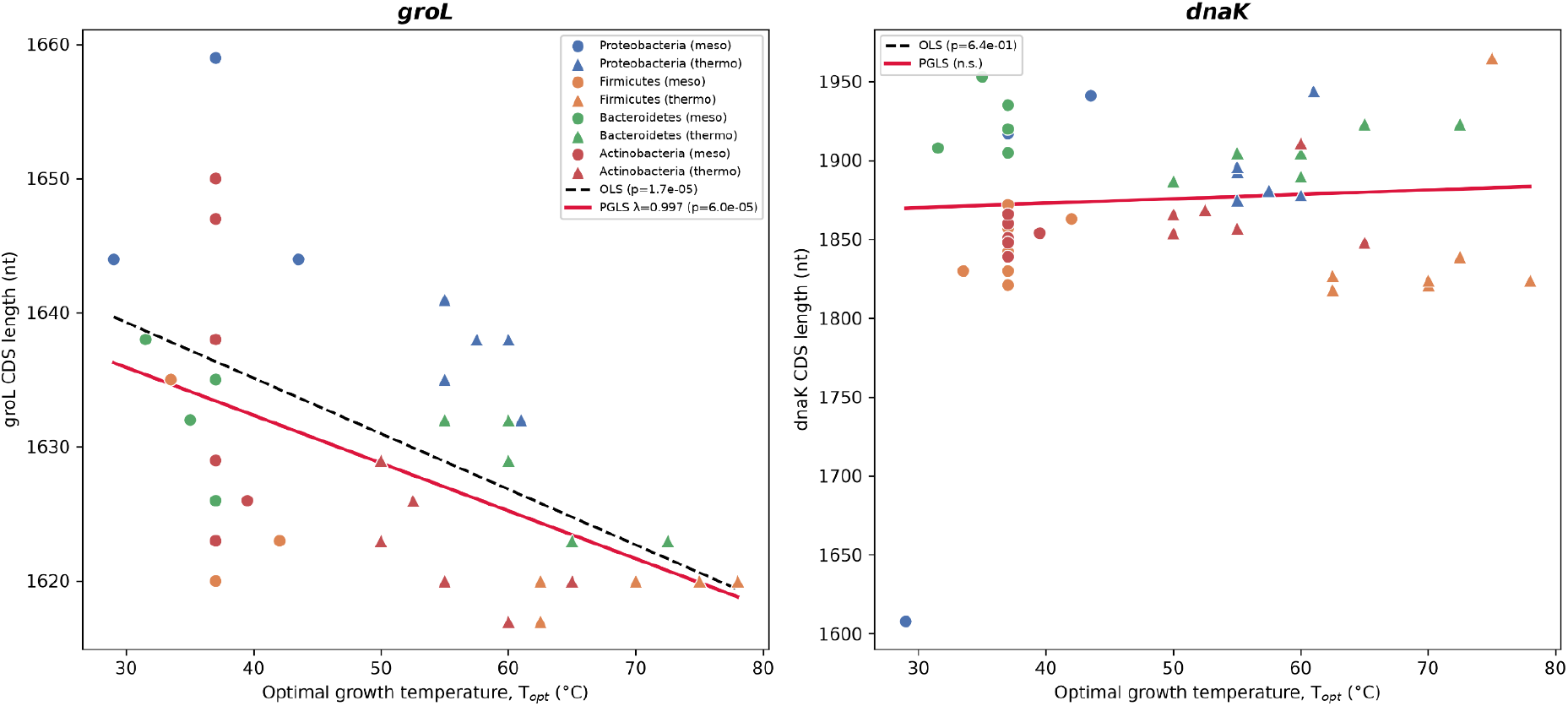
Coding sequence length against optimal growth temperature for *groL* (left) and *dnaK* (right). Points are coloured by phylum; circles denote mesophiles and triangles thermophiles. Dashed black line, ordinary least squares fit; solid red line, phylogenetic generalised least squares fit with Pagel’s λ estimated by maximum likelihood from the 16S tree. The *groL* relationship is significant under both models; *dnaK* under neither.

### The length relationship survives phylogenetic correction

After matching tip labels between the 16S tree and the phenotype table, all the 50 strains were retained. Ordinary least squares regression of groL length on T_opt_ on this matched set gives a slope of -0.415 nt °C^-1^ (P = 1.7 × 10^-5^). Under phylogenetic generalised least squares with Pagel’s λ estimated by maximum likelihood, the slope is -0.356 nt °C^-1^ with P = 6.0 × 10^-5^ (Figure 2). The maximum-likelihood estimate was λ = 0.997. A λ near unity indicates substantial phylogenetic structure in the model residuals, which is precisely the condition under which uncorrected regression would be untrustworthy; that the relationship persists after correction indicates it is not an artefact of shared ancestry. The corresponding dnaK model remains non-significant.

### The length difference is confined to the disordered C-terminal tail

Aligning all 50 GroEL proteins and comparing per-column gap frequencies between thermal groups localises the signal sharply (Figure 3). Across residues 1-523 of the *E. coli* reference, spanning the equatorial, intermediate and apical domains that constitute the folded chaperonin body, excess deletion in thermophiles is negligible and unsystematic. Approximately three-quarters of the total positive excess-gap mass (3.00 of 4.08 column-equivalents, 73.5%) falls within the last 30 alignment columns, which span *E. coli* residues 524-548 together with three insertion columns internal to that region. This partition is insensitive to the boundary chosen: any cut-off between the last 30 and last 50 columns yields the identical 73.5%, since no positive excess mass occurs in columns 519-539; consistent with this, 12 of the 13 alignment columns showing more than a 15 percentage-point excess gap rate in thermophiles lie within this terminal region. The largest single-column excess (0.36) occurs at *E. coli* residue 536. As a negative control, DnaK protein length showed no significant difference between mesophiles and thermophiles (mean 623.6 ± 23.1 aa versus 624.6 ± 13.3 aa; Mann-Whitney P = 0.71, Cliff’s δ = +0.06), indicating that the length reduction is specific to GroEL rather than a general feature of chaperone proteins. This terminal ∼25-residue segment corresponds to the intrinsically disordered, glycine/methionine-rich C-terminal extension of GroEL, which terminates in the conserved (GGM)_4_M motif and is not resolved in crystal or cryo-EM structures of the chaperonin^16^,^17^.

**Figure 3.**
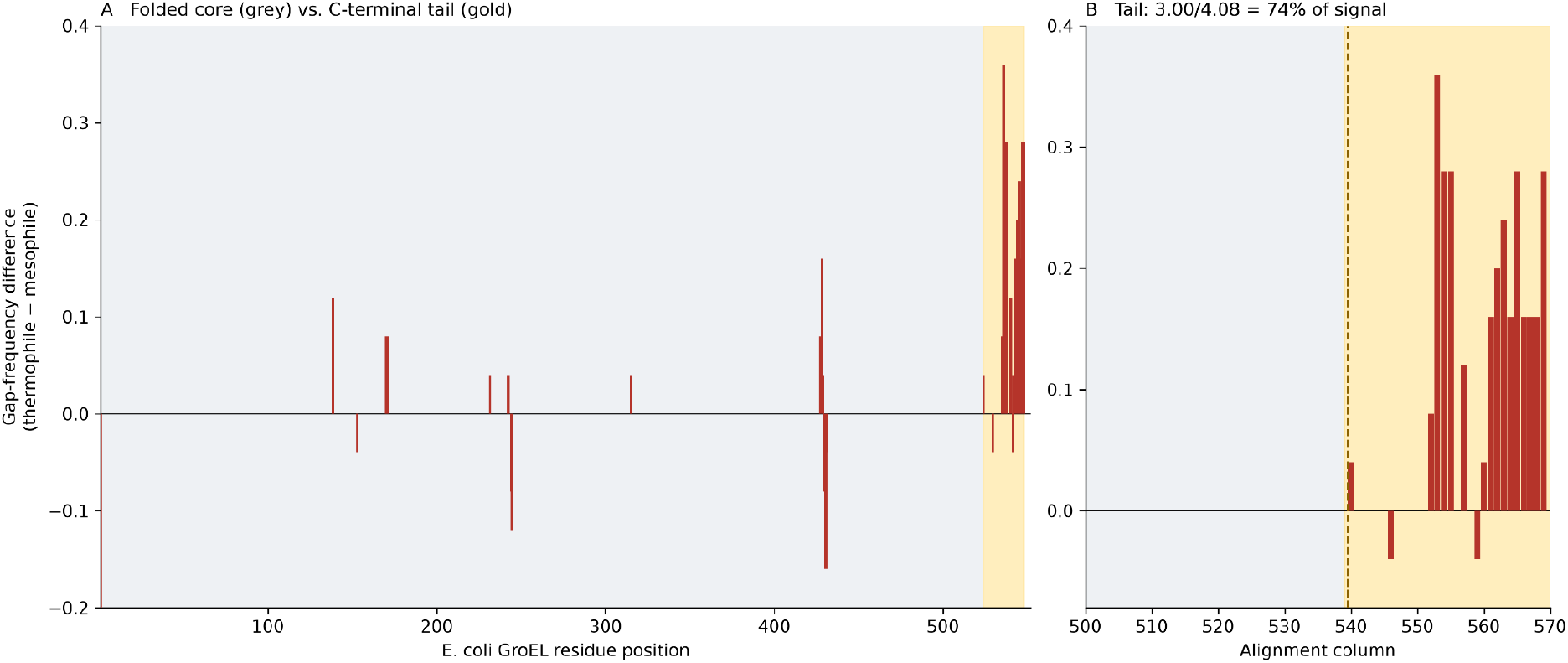
Per-column difference in gap frequency (thermophile - mesophile) across the GroEL protein alignment, mapped to *E. coli* GroEL residue numbering. Positive values indicate residues more often absent in thermophiles. The shaded terminal region marks the intrinsically disordered glycine/methionine-rich C-terminal tail, which is not resolved in structural studies. Excess deletion is negligible across the folded core (residues 1-523) and concentrated in the tail.

**Table 2.** Protein lengths of GroEL (full and C-terminal tail) and DnaK (negative control).

| **Protein region** | **Mesophile mean ± SD** | **Thermophile mean ± SD** | **Mann-Whitney P** | **Cliff’s δ** |
|---|---|---|---|---|
| GroEL full length | 544.4 ± 3.3 aa | 541.2 ± 2.4 aa | 0.0005 | +0.57 |
| GroEL C-terminal tail | 21.8 ± 3.3 aa | 18.9 ± 2.3 aa | 0.0017 | +0.52 |
| DnaK full length (control) | 623.6 ± 23.1 aa | 624.6 ± 13.3 aa | 0.71 | +0.06 |

## Discussion

Two principal findings emerge.

The first concerns mRNA secondary structure. Across fifteen thermodynamic and structural descriptors, two folding temperatures and two genes, the raw folding metrics did not distinguish thermophilic from mesophilic bacteria. This held for the heat-shock chaperone transcripts themselves, arguably the most favourable case one could construct for detecting thermal tuning of mRNA folding. In that limited sense the result is consistent with Jegousse *et al*.’s^1^ interpretation that bacterial mRNA is compositionally biased toward flexibility rather than toward niche-appropriate structure. However, when dinucleotide composition was controlled by Altschul-Erickson shuffling, a residual structural signal became detectable: thermophile sequences, particularly *groL*, yielded significantly more negative structural z-scores than mesophile sequences, especially at the higher folding temperature. The residual signal is modest in magnitude and weaker in *dnaK*, but it is reproducible across different numbers of shuffles. Thus the absence of a difference in raw free-energy measures does not imply the complete absence of composition-independent structural organisation.

The second finding is positive and concerns protein architecture rather than RNA. The only descriptor that cleanly separates the groups at the level of raw sequence properties is coding-sequence length, and the difference localises to a disordered peptide extension rather than to the folded core. Shortening of surface loops and terminal extensions is an established route to protein thermostability^18^, and the GroEL C-terminal tail is a plausible target: it is reported to contribute to client folding, to form a barrier limiting client escape from the base of the chaperonin barrel, and to adopt polyproline II helical conformation rather than being wholly unstructured^16^. Its functional status has been revised over time; an early study concluded the conserved glycine-glycine-methionine terminus was dispensable^19^, whereas more recent sequence and viability analyses argue that the disordered segment is essential and that bacteria carrying multiple chaperonin paralogues consistently retain at least one bearing the glycine/methionine-rich terminus^17^.

We emphasise that we have not tested any functional consequence of tail shortening. Experimental work nevertheless supports a temperature-related role for this region: swapping the C-terminal segment of the GroEL equatorial domain between species with different optimal growth temperatures substantially altered the active temperature range of the resulting chimeras^20^. In archaeal group II chaperonins the C-terminal residues likewise contribute to thermostability^21^. One possibility, therefore, is that a shorter disordered tail alters encapsulation geometry or client-release kinetics in a manner favoured at elevated temperature, consistent with the broader pattern of loop and terminal truncation as a route to protein thermostability^18^. An alternative is that the tail is simply a region of relaxed constraint where indel variation accumulates, and that its correlation with T_opt_ reflects a correlate of thermophily other than direct selection on tail length. Distinguishing these interpretations would require functional assays or explicit tests for selection, neither of which we performed. The magnitude involved, roughly three codons on a gene of ∼1630 nt, is small in absolute terms even though the group separation is statistically clear.

Several limitations bound these conclusions. With 25 strains per group we are adequately powered for effects of the size actually detected (δ ≈ 0.5) but not for small ones, so undetected effects should be described as not detected at this sample size rather than as absent. Our tree is a single-gene 16S phylogeny with substitution-based branch lengths, not a multilocus chronogram. The cohort is bacterial and excludes hyperthermophiles by design, so extrapolation above 80°C is unwarranted. Finally, we analysed complete coding sequences because regulatory RNA thermometers in bacteria typically reside in 5′ untranslated regions rather than within coding sequence, a targeted analysis of 5′ regions and their upstream flanks remains an obvious and untested extension of this work.

## Supporting information

16S rRNA phylogenetic tree of the 50 bacterial strains

