## Supplementary material for "Chaperone mRNA secondary structure does not strongly track thermal adaptation in bacteria, but *groL* coding sequence shortens in its disordered C-terminal tail": 16S rRNA phylogenetic tree of the 50 bacterial strains

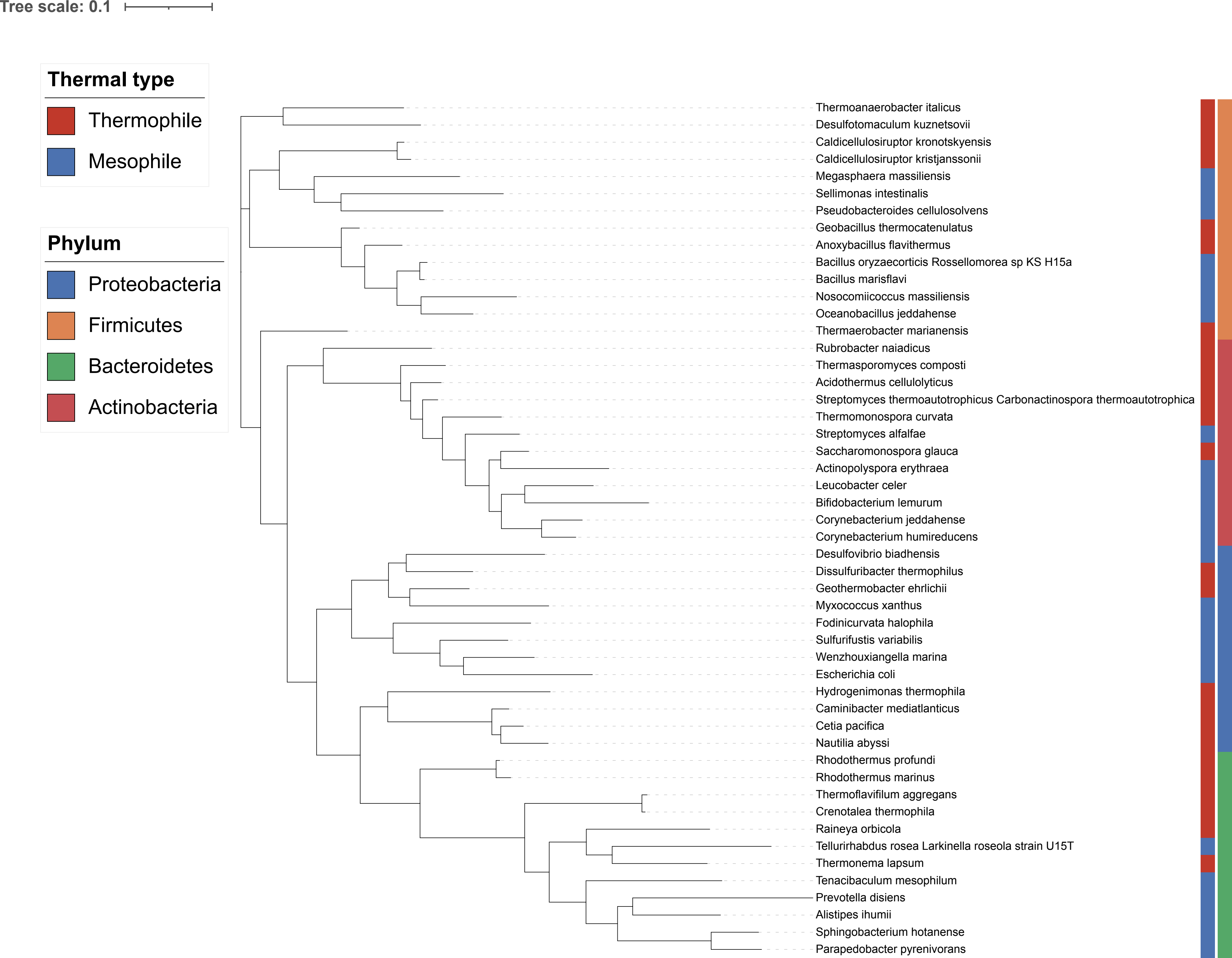

Tree scale: 0.1
Thermal type
Thermophile
Mesophile
Phylum
Proteobacteria
Firmicutes
Bacteroidetes
Actinobacteria
Thermoanaerobacter italicus
Desulfotomaculum kuznetsovii
Caldicellulosiruptor kronotskyensis
Caldicellulosiruptor kristjanssonii
Megasphaera massiliensis
Sellimonas intestinalis
Pseudobacteroides cellulosolvens
Geobacillus thermocatenulatus
Anoxybacillus flavithermus
Bacillus oryzaecorticis Rossellomorea sp KS H15a
Bacillus marisflavi
Nosocomiicoccus massiliensis
Oceanobacillus jeddahense
Thermaerobacter marianensis
Rubrobacter naiadicus
Thermasporomyces composti
Acidothermus cellulolyticus
Streptomyces thermoautotrophicus Carbonactinospora thermoautotrophica
Thermomonospora curvata
Streptomyces alfalfae
Saccharomonospora glauca
Actinopolyspora erythraea
Leucobacter celer
Bifidobacterium lemurum
Corynebacterium jeddahense
Corynebacterium humireducens
Desulfovibrio biadhensis
Dissulfuribacter thermophilus
Geothermobacter ehrlichii
Myxococcus xanthus
Fodinicurvata halophila
Sulfurifustis variabilis
Wenzhouxiangella marina
Escherichia coli
Hydrogenimonas thermophila
Caminibacter mediatlanticus
Cetia pacifica
Nautilia abyssi
Rhodothermus profundi
Rhodothermus marinus
Thermoflavifilum aggregans
Crenotalea thermophila
Raineya orbicola
Tellurirhabdus rosea Larkinella roseola strain U15T
Thermonema lapsum
Tenacibaculum mesophilum
Prevotella disiens
Alistipes ihumii
Sphingobacterium hotanense
Parapedobacter pyrenivorans
